# METEOR: joint genome-scale reconstruction and enzyme prediction

**DOI:** 10.64898/2026.02.01.701942

**Authors:** Kexin Niu, Maxat Kulmanov, Robert Hoehndorf

## Abstract

**Motivation:** Current machine learning methods for enzyme function prediction primarily treat proteins as independent entities, ignoring the metabolic context in which they operate. This reductionist approach often generates biologically implausible annotations that fail to satisfy stoichiometric or thermodynamic constraints. While genome-scale metabolic models (GEMs) enforce systemic coherence, traditional reconstruction workflows decouple functional annotation from network assembly, resulting in information loss during the discretization of enzymatic evidence.

**Results:** We developed METEOR, a framework that integrates continuous confidence scores from deep learning models directly into a Mixed-Integer Linear Programming (MILP) formulation. By conditioning predictions on global metabolic requirements, METEOR resolves functional ambiguities where sequence signals are weak. Evaluation on the Price-149 dataset shows that our method yields physiologically viable networks capable of sustaining growth and improve protein-level predictions over baseline methods. Validation on 6,894 bacterial genomes reveals that system-aware refinement increases the recall of experimentally observed phenotypes substantially. Furthermore, we show that constraint-driven selection can still improve annotation performance in highly incomplete genomes. Our results suggest that functional annotation should be treated as a unified inference problem where global system constraints supervise local predictions.

**Availability:** METEOR is freely available at https://github.com/bio-ontology-research-group/Meteor

**Supplementary information:** Supplementary data are available at Bioinformatics online.

## Introduction

The accurate annotation of enzymatic activities within newly sequenced genomes constitutes a fundamental challenge in computational biology. Rapid advances in high-throughput sequencing technologies have facilitated the generation of massive genomic datasets, necessitating the development of automated functional prediction tools (Kustatscher *et al*., 2022; Kumar *et al*., 2024). However, functional annotation lags behind data generation. Various computational methods predict functions from different protein features (You *et al*., 2018; Kulmanov and Hoehndorf, 2020; Cao and Shen, 2021; Lai and Xu, 2022; Boadu *et al*., 2023; Gligorijević *et al*., 2021). Computational methods for protein function annotation have evolved significantly, moving from alignment-based approaches (Desai *et al*., 2011; Altschul *et al*., 1990, 1997) to machine learning architectures (Kulmanov and Hoehndorf, 2020; Cao and Shen, 2021; Gligorijević *et al*., 2021; Lai and Xu, 2022). Early methods primarily used homology detection (Bateman *et al*., 2004; Cantalapiedra *et al*., 2021), inferring function by transferring annotations from characterized sequences to uncharacterized homologs. However, recent developments use deep learning to capture complex patterns within protein sequences and structures (Cao and Shen, 2021; Wang *et al*., 2025; Kulmanov and Hoehndorf, 2020; Yuan *et al*., 2023). Several methods specifically predict enzymes, usually expressed using the Enzyme Commission (EC) numbers. DeepEC (Ryu *et al*., 2019) employs convolutional neural networks (CNNs) to hierarchically classify enzymatic activities. More recent models, including DeepProZyme (Kim *et al*., 2023) and EnzBert (Buton *et al*., 2023), integrate transformer-based architectures and attention mechanisms to process protein sequences, expanding the range of predictable classes to include translocases. Additionally, contrastive learning frameworks like CLEAN (Yu *et al*., 2023) maximize the separation between protein embeddings and negative samples to enhance representation quality.

Although these methods have evolved, they predict functions by treating proteins as independent entities, ignoring genomic context. Specifically, they infer function from local (i.e., protein-specific) features, disregarding the systemic context in which proteins operate (Yu *et al*., 2023; Ryu *et al*., 2019; Kim *et al*., 2023; Buton *et al*., 2023). Nevertheless, biological systems impose strict constraints on enzymatic activity: proteins do not function in isolation but act as components of integrated biochemical networks subject to stoichiometric and thermodynamic limitations (He *et al*., 2016; Orth *et al*., 2010). Due to these constraints, two proteins may have different enzymatic functions in different organisms, even if they have 100% sequence identity. Prediction algorithms that ignore the interdependencies that exist in biological systems may generate biologically implausible annotations (Tawfiq *et al*., 2025).

Genome-Scale Metabolic Models (GEMs) explicitly model biological systems (Chen *et al*., 2024; He *et al*., 2016). GEMs formalize and represent the biochemical reaction networks of an organism, linking genotype to phenotype through gene–protein– reaction (GPR) associations (Di Filippo *et al*., 2021). While these genome-scale models enforce system-wide constraints, such as mass balance and pathway completeness, conventional reconstruction workflows treat functional annotation and model building as two different and separated processes (Jenior *et al*., 2023; Machado *et al*., 2018). Methods that generate metabolic models usually use binary presence/absence predictions to generate a draft model, and subsequently apply gap-filling algorithms to restore connectivity (Jenior *et al*., 2023; Machado *et al*., 2018; Thiele *et al*., 2014).

Standard gap-filling methodologies have evolved significantly but still constrained by their underlying optimization logic. Early parsimony-based methods, like fastGapFill (Thiele *et al*., 2014) and the more recent COBRApy-integrated tool Reconstructor (Jenior *et al*., 2023), search for the minimum number of reactions from a universal database to restore network connectivity. While these tools employ coarse-grained category weights—such as penalizing transport reactions to prioritize internal metabolic flux—they fundamentally prioritize mathematical compactness over biological likelihood, often yielding “shortcuts” that lack physiological relevance. To incorporate biological evidence, subsequent tools have introduced genomic-weighted objectives. For instance, gapseq (Zimmermann *et al*., 2021) utilizes sequence alignment score (bitscores) mapped to penalty weights via a heuristic step function. However, this discretization leads to a degradation of information resolution, discrete penalty tiers. Even top-down approaches like CarveMe (Machado *et al*., 2018), which emoploy Mixed-Integer Linear Programming (MILP) to extract organism-specific models from a universal template, rely on binary selection strategies during the draft construction phase. These disjointed, two-step paradigm strategies are prone to irreversible inclusion of isolated enzymes—where a reaction is included based on high local sequence confidence despite its broader metabolic pathway lacking genomic support. Subsequently, gap-filling algorithms will add multiple unsubstantiated reactions to bridge the gaps. By thresholding functional annotations into presence/absence calls before network integration, these workflows become susceptible to error propagation. Consequently, current paradigms fail to achieve Full-scale Continuous Score Optimization, leading to significant biological information loss during the discretization of enzymatic evidence.

Integrating genome-scale constraints directly into the enzyme prediction process presents an opportunity to improve enzyme prediction accuracy. By conditioning enzyme prediction probabilities on metabolic feasibility, systems biology methods can resolve ambiguities where sequence-based signals are weak. For instance, the presence of a complete metabolic pathway required for biomass production provides strong systemic evidence for the existence of a missing enzyme, even if sequence similarity or prediction scores are marginal. Therefore, we reason the an approach to address protein (enzyme) function prediction from a systems biology perspective enables the prediction of enzymatic activities that are both genomically supported and metabolically coherent.

We developed METEOR, a theoretical and computational framework that uses genome-scale metabolic constraints to improve enzymatic function predictions. We developed an optimization-based reconstruction method that integrates continuous probabilistic scores from deep learning models directly into a Mixed-Integer Linear Programming (MILP) formulation of the metabolic network. Rather than thresholding predictions into binary values, our approach forces the reconstructed network to satisfy global metabolic requirements, such as stoichiometry and biomass production, while simultaneously minimizing uncertainty added to the original ML-derived confidence scores. This approach enables the construction of functional annotations that optimize both for genomic evidence and biological system constraints.

## Materials and methods

### Data preparation

#### Dataset and reference genomes

We used the Price-149 dataset (Price *et al*., 2018) as the primary dataset for evaluating the protein-level accuracy of enzyme prediction. The Price-149 dataset is also widely used to evaluate and compare the performance of enzyme function prediction methods (Yu *et al*., 2023; Kim *et al*., 2023; Song *et al*., 2024). The Price-149 dataset consists of 149 experimentally validated proteins with their EC numbers. The original study (Price *et al*., 2018) validated over 400 enzymes on 32 genomes; Price-149 dataset contains 149 enzymes present in 22 genomes (Yu *et al*., 2023). We obtained the genome assemblies for all 22 genomes from which the Price-149 dataset proteins are derived. Detailed see Table S1.

#### Sequence-based prediction and confidence scoring

We used several state-of-the-art enzyme function prediction methods for validation and comparison, and to provide prediction scores which we used as input probabilities for protein–enzyme–reaction associations.

We used DeepEC (Ryu *et al*., 2019), which employs three independent convolutional neural networks (CNNs) to hierarchically predict Enzyme Commission (EC) numbers. The first CNN performs binary classification of the protein sequence, while subsequent networks classify the enzyme into specific subclasses and sub-subclasses.

We also used DeepProZyme (Kim *et al*., 2023), which uses a transformer-based architecture to predict enzymatic function. Building upon DeepEC, DeepProZyme extends prediction capabilities to include translocases (EC class 7), increasing the total number of supported EC classes to 5,360.

We also used EnzBert (Buton *et al*., 2023), a model based on protein language models. The EnzBert model incorporates a transformer architecture with self-attention mechanisms to capture dependencies within protein sequences. EnzBert initializes weights from a pre-trained ProtBert-BFD model (Elnaggar *et al*., 2021) and is fine-tuned specifically for EC class prediction tasks.

### System-aware refinement via metabolic optimization

We show the overall pipeline of the METEOR framework in Figure 1. This pipeline integrates protein-level deep learning predictions with global metabolic constraints through Mixed-Integer Linear Programming (MILP) to ensure that functional annotations remain metabolically coherent. Consequently, this system-aware refinement helps resolve cases where sequence signals remain weak to improve the physiological viability of the resulting metabolic models.

**Figure 1.**
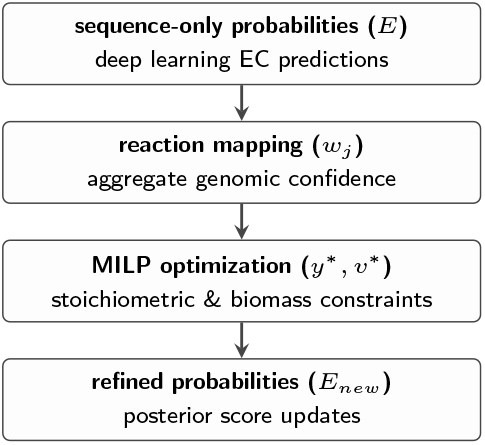
The METEOR framework uses a sequential refinement pipeline to resolve functional ambiguities by anchoring protein-level predictions to global metabolic requirements.

#### Formulation of reaction costs from prediction confidence

As shown in the overview of our method (Figure 1), METEOR starts by processing sequence-only probabilities (*E*). To bridge sequence-based enzyme predictions with metabolic modeling, we define a framework linking proteins, functional assignments, and biochemical reactions. Let 𝒫 denote the set of proteins in the target genome, ℰ 𝒞, the set of all Enzyme Commission (EC) numbers represented in the scoring matrix, and ℛ the set of reactions in a universal metabolic database. We used a protein-centric predictor to generate a confidence matrix **E** ∈ [0, 1]^|𝒫|*×*|ℰ 𝒞|^, where *E*_*p,e*_ represents the probability of protein *p* being assigned to EC number *e*. These functional assignments are mapped to biochemical reactions via a curated mapping ℳ : ℰ 𝒞 → ℛ. For each reaction *j* ∈ ℛ, we determined the aggregate genomic confidence *w*_*j*_ by the highest-scoring protein-EC pair among all possible assignments that map to that specific reaction, formulated as

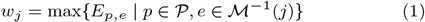

Using a predefined confidence threshold *θ* ∈ [0, 1], we established an initial reaction pattern 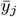, where 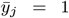 and 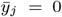 otherwise, providing a prediction-informed foundation for the subsequent optimization process.

### Integration of metabolic tasks and costs

We defined modification costs that penalize deviations from the genomic evidence to ensure the resulting metabolic model remains biologically grounded. Let

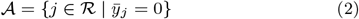

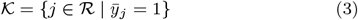

represent the sets of reactions considered for addition or removal, respectively. The addition cost 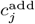 for *j* ∈ 𝒜 penalizes the inclusion of reactions lacking genomic support, defined as:

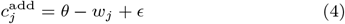

where *ϵ* is a small regularization constant ensuring parsimonious selection. For all experiments in this paper, we use default parameters with *ϵ* = 1*e* − 3 (Figure S5). Conversely, the removal cost 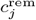 for *j* ∈ 𝒦 penalizes the exclusion of reactions strongly supported by sequence data. This cost aggregates evidence from all protein-EC pairs mapping to reaction *j* that exceed the threshold, calculated as:

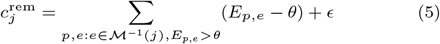

These costs transformed network reconstruction into an evidence-weighted optimization problem, balancing enzymatic plausibility with the requirement to fill functional metabolic gaps.

### The Mixed-Integer Linear Programming (MILP) model

The metabolic network structure is represented by a stoichiometric matrix **S** ∈ ℝ^*M ×*|*ℛ*|^ for *M* metabolites. Let **v** ∈ ℝ ^|*ℛ*|^ be the steady-state flux vector, subject to biophysical realistic bounds *lb*_*j*_ and *ub*_*j*_ for each reaction *j* ∈ ℛ. The selection of reactions for the final organism-specific model is represented by a binary vector **y** ∈ {0, 1}^|*ℛ*|^.

We formulated the system-aware refinement of ML scores as a Mixed-Integer Linear Programming (MILP) problem using stoichiometric necessity and growth constraint. The objective finds the optimal vector **y** and flux vector **v** that minimize the weighted cost from the initial predictions while enforcing mandatory metabolic growth (e.g., biomass production):

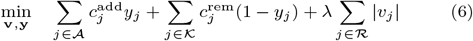

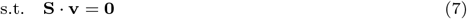

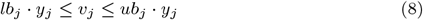

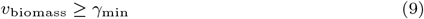

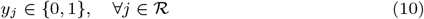

Equation 7 enforces the steady-state mass balance. Equation 8 links reaction selection *y*_*j*_ to flux *v*_*j*_. Equation 9 ensures the model satisfies a mandatory biomass flux requirement *γ*_min_. A secondary objective term *λ* ∑ |*v*_*j*_ | (with *λ* ≪ 1) favors parsimonious flux distributions. For all experiments in this paper, we use default parameters with *γ*_min_ = 1.0, *λ* = 1*e* − 6 (Figure S5).

### Refinement of enzyme predictions based on metabolic solution

The optimal solution **y**^∗^ derived from the Mixed-Integer Linear Programming (MILP) formulation enables a posteriori refinement of the original confidence matrix **E** by back-propagating system-level constraints. Based on the selected reactions where 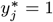, we define the set of active enzymes

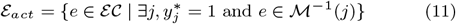

and the set of muted enzymes

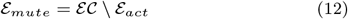

For activation, if an EC in ℰ _*act*_ was not previously predicted, (i.e., its initial maximum score in the genome was below the threshold *θ*) we identify the top proteins with the highest latent scores (or those matching the 3-digit EC prefix) and update their scores to 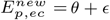 previously predicted, the score of the most probable protein is boosted. For muted enzymes in ℰ _*mute*_, we suppress the scores of all associated proteins by setting an upper bound of 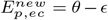 If the initial maximum score for an EC within the genome is zero, we skip this EC avoid giving random assignment.

### Implementation and performance assessment

#### Computational implementation and solvers

We implemented our framework in Python (3.10) using the PuLP (v3.2.2) library (Mitchell *et al*., 2011) as the modeling interface. We formulated the reconstruction problem as a Mixed-Integer Linear Programming (MILP) task, enabling the simultaneous optimization of discrete reaction selections (binary variables) and continuous metabolic fluxes (continuous variables). We solved the resulting MILP problems using the COIN-OR Branch-and-Cut (CBC) solver (Forrest *et al*., 2024).

### Evaluation metrics

To evaluate our method’s performance, we used standard classification metrics, including F1 score, recall, precision, and the Area Under the Receiver Operating Characteristic Curve (ROC-AUC) (Zhou *et al*., 2014). We defined True Positives (TP) as correctly identified enzymatic functions, False Positives (FP) as incorrect assignments, and False Negatives (FN) as missed annotations. We calculated the primary metrics as follows:

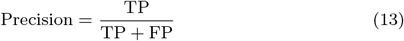

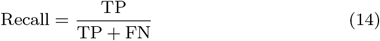

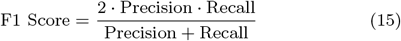

We also reported the ROC-AUC (Zhou *et al*., 2014), integrating the True Positive Rate (TPR) as a function of the False Positive Rate (FPR) across all threshold settings:

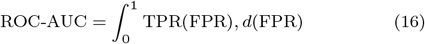

We used Scikit-learn package (Pedregosa *et al*., 2011) to calculate the F1-score, recall, precision, and ROC-AUC for all experiments in this study.

#### Model quality validation with MEMOTE

To evaluate the quality of the reconstructed genome-scale metabolic models (GEMs), we utilized the standardized benchmarking tool MEMOTE (v0.17.0) (Lieven *et al*., 2020). This open-source platform provides a systematic framework for quantifying model quality across several core dimensions, including stoichiometry, mass and charge balances, and metabolic connectivity.

Notably, MEMOTE employed Flux Balance Analysis (FBA) to assess the growth potential of the model. FBA is a constraint-based mathematical approach that predicts the flow of metabolites through a biochemical network by assuming a pseudo-steady state. FBA leverages stoichiometric constraints derived from reaction networks and predicts feasible metabolic flux distributions by solving a linear optimization problem. Formally, FBA is formulated as a linear programming problem:

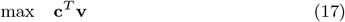

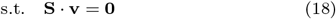

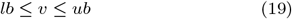

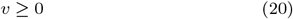

where **c** defines the objective function, typically corresponding to biomass production, energy generation, or the synthesis of specific metabolites.

We use default media with rich medium for all metabolic models generated in this study.

## Results

### Systemic constraints boost enzymatic functional annotations

We first evaluated the internal logic of METEOR by evaluating the capacity of genome-scale metabolic constraints to refine enzymatic function predictions by integrating systemic biological requirements into the annotation pipeline. Traditional sequence-only classifiers, such as DeepEC (Ryu *et al*., 2019), EnzBert (Buton *et al*., 2023), and DeepProZyme (Kim *et al*., 2023), predict functions for proteins as independent entities. However, enzymes must operate within a coherent biochemical network to sustain biomass production.

To establish a baseline for protein-level accuracy, we used the Price-149 dataset, which consists of 149 experimentally validated enzymes (Sanderson *et al*., 2023; Price *et al*., 2018). We mapped these proteins to their original 22 bacterial species to ensure the systemic context matched the genomic evidence.

Table 1 demonstrates that our framework consistently improved predictive performance across all architectures. For instance, the optimization increased the F1-score of DeepProZyme from 0.434 to 0.463 and improved its ROC-AUC from 0.697 to 0.710. These results indicate that conditioning predictions on metabolic feasibility effectively filters prediction noise, enabling the system to resolve cases where sequence-based signals are ambiguous.

**Table 1.** Comparative evaluation of enzyme prediction accuracy on the Price-149 dataset.

| Methods |  | Precision | Recall | F1 | AUC |
| --- | --- | --- | --- | --- | --- |
| EnzBert | Sequence-only | 0.385 | 0.349 | 0.343 | 0.673 |
|  | System-aware | 0.424 | 0.355 | 0.354 | 0.677 |
| DeepEC | Sequence-only | 0.263 | 0.132 | 0.158 | 0.565 |
|  | System-aware | 0.289 | 0.138 | 0.168 | 0.568 |
| DeepProZyme | Sequence-only | 0.584 | 0.395 | 0.434 | 0.697 |
|  | System-aware | **0.613** | **0.421** | **0.463** | **0.710** |

**Sequence-only** indicates the performance of ML-derived scores. **System-aware** indicates the performance of refinement scores through genome-scale metabolic constraints. Bold values highlighting the superior performance in each category.

### Evaluation of model functionality and physiological viability

Beyond individual protein annotations, we evaluated whether these refinements yield improved system-level performance in the resulting metabolic models. We compared our models against the Reconstructor pipeline (Jenior *et al*., 2023), which serves as a traditional baseline for automated reconstruction. In traditional workflows, a draft model is built from high-confidence sequence matches, and a gapfilled (GF) model is subsequently produced by adding the minimum number of reactions required to bridge pathway gaps based on topological heuristics (Lewis *et al*., 2010).

We employed the MEMOTE benchmarking suite to quantify model quality (Table 2). A fundamental requirement for any biological model is adherence to physical laws, measured by the *Consistency* score. All models generated by our framework exhibited high self-consistency, with the DeepEC-based reconstruction achieving a score of 0.893. While the Reconstructor (GF) baseline achieved a higher *Total* score, this was driven by superior Systems Biology Ontology (SBO) (Courtot *et al*., 2011) coverage (0.727). SBO scores measure standardized semantic labeling rather than biochemical logic or network connectivity (Ravikrishnan and Raman, 2015; Lieven *et al*., 2020).

**Table 2.** Comparison of metabolic consistency and functionality metrics.

| Methods | Consistency |  | Model Functionality |  |  |  |
| --- | --- | --- | --- | --- | --- | --- |
|  | Score | SBO | Growth<sup>1</sup> | Gaps<sup>2</sup> | Blk.%<sup>3</sup> | Unreal.<sup>4</sup> |
| EnzBert | 0.880 | 0.636 | 12.65 | 1.09 | 0.53 | 3.77 |
| DeepEC | **0.893** | 0.636 | **12.66** | **0.64** | 0.54 | 3.91 |
| DeepProZyme | 0.877 | 0.636 | 8.56 | 1.36 | **0.48** | 3.77 |
| Reconstructor (Draft) | 0.863 | 0.545 | 0.00 | 0.00 | 0.67 | **0.00** |
| Reconstructor (GF) | 0.869 | **0.727** | 0.00 | 5.00 | 0.60 | **0.00** |

Bold values highlighting the superior performance in each category.
<sup>1</sup>Growth indicates predicted growth rate ( $h^{-1}$ ).
<sup>2</sup>Gaps indicates blocked biomass precursors.
<sup>3</sup>Blk.% indicates fraction of blocked reactions.
<sup>4</sup>Unreal indicates Unrealistic Growth Rate tests

Our models also significantly reduced gaps in biomass synthesis, achieving a minimum of 0.64 blocked precursors in the DeepEC-based optimization, whereas the Reconstructor (GF) baseline contained blocked precursors on average. Consequently, all models produced by our framework achieved biologically relevant biomass production rates (8.56 to 12.66*h*^−1^), while the baseline models remained physiologically inactive (0.00*h*^−1^) under default medium conditions (Figure S1).

### Systemic redundancy provides resilience against genome incompleteness

To evaluate whether METEOR functions in practical scenarios involving partial genomes, such as metagenome-assembled genomes (MAGs), we benchmarked performance across simulated levels of genome incompleteness. We used the Price-149 dataset and its 22 original bacterial genomes to establish a test set. We simulated information loss by randomly removing proteins from the genomic inputs at intervals ranging from 10% to 99%.

Figure 2 depicts the mean performance metrics across these trials. The performance of METEOR remains stable until a significant fraction of the genome is removed. Even with 90% of the genome proteins removed, METEOR provided a ROC-AUC improvement over the baseline from 0.697 to 0.713. We attribute this resilience to constraint-driven selection. As the pool of available proteins shrinks, the MILP optimization is forced to rely on the remaining candidates to satisfy the mandatory biomass objective. Because the test proteins, many of which represent essential core metabolic functions, are retained, the metabolic constraints act as a selection pressure that forces their inclusion in the model to bridge pathway gaps. Simultaneously, the removal of non-essential “background” proteins reduces the solution space, effectively eliminating potential false positives (distractors) that could otherwise lead to erroneous assignments. Therefore, in highly fragmented genomes, the requirement for physiological viability forces the algorithm to correctly identify and employ the few available essential enzymes, maintaining high precision and recall despite the loss of broader context (Figure S2).

**Figure 2.**
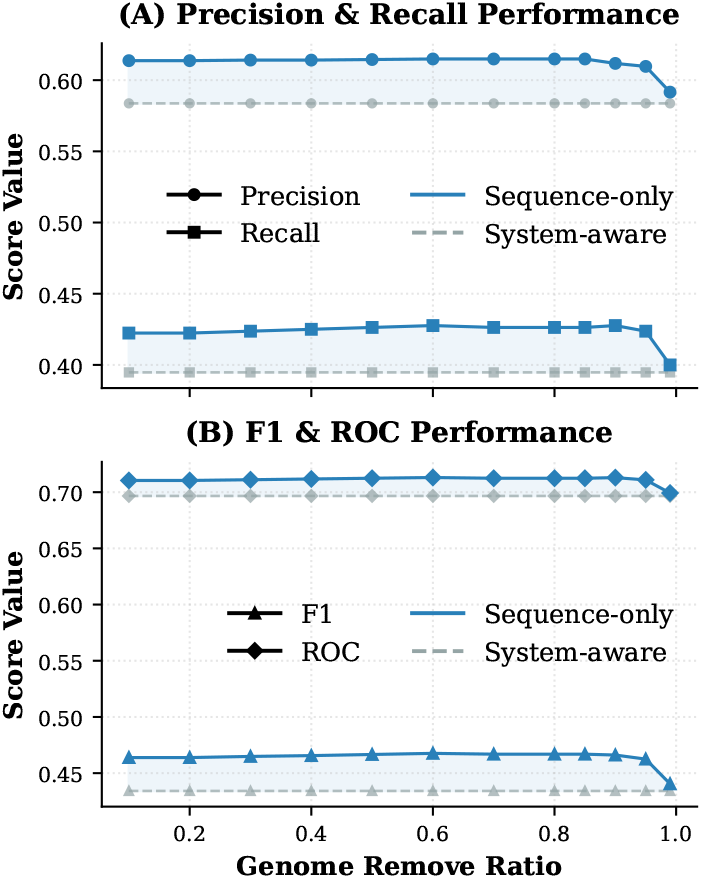
Genome-scale optimization with incomplete genome on the Price-149 dataset. Data points represent the mean F1-score across five trials. METEOR maintained stable performance until 90% genome removal.

### Phenotype validation through system-level realized enzymatic activities

To validate the physiological accuracy of the models generated by METEOR, we benchmarked their ability to capture experimentally observed phenotypes. We queried the BacDive database (Schober *et al*., 2025) to extract a dataset comprising 6,894 genomes and 91,633 distinct phenotypic entries. Our analysis focused on experimental enzyme activity assays to verify whether the computational models correctly retained enzymes known to be active *in vivo*.

We annotated the complete genomes using METEOR to generate optimized metabolic networks. To ensure that the reconstruction was not artificially limited by environmental scarcity, we simulated a “complete” medium condition, allowing the uptake of all library nutrients. This configuration maximizes the theoretical metabolic potential, ensuring that any enzyme capable of contributing to biomass production is eligible for inclusion. We then cross-referenced the resulting models against the BacDive records to determine if the experimentally validated enzymes were present in the final, system-aware reconstructions.

Table 3 summarizes the performance across the ten most frequent Enzyme Commission (EC) categories. METEOR demonstrated superior alignment with realized phenotypes, achieving an overall recall of 0.751 and an F1-score of 0.654, compared to a recall of 0.720 and F1-score of 0.648 for the sequence-only baseline. We observed a small decrease in precision, which dropped from 0.589 to 0.580. This inverse trend suggests that while the optimization framework effectively recovers essential enzymes missed by sequence classifiers, it also recruits proteins to satisfy biomass constraints that are not currently recorded in the phenotype database. Given that experimental annotations are often non-exhaustive, a portion of these “false positives” may also represent true, biologically required enzymatic activities that are simply absent from the reference dataset.

**Table 3.** Comparative performance analysis across the ten most frequent Enzyme Commission (EC) categories in the BacDive reference dataset.

| Category |  | Entries | Precision | Recall | F1-score |
| --- | --- | --- | --- | --- | --- |
| Overall | Sequence-only | 91,633 | **0.589** | 0.720 | 0.648 |
|  | System-aware | 91,633 | 0.580 | **0.751** | **0.654** |
| EC 1.11.1.6 | Sequence-only | 6,108 | 0.900 | 0.884 | 0.892 |
|  | System-aware | 6,108 | 0.900 | **0.886** | **0.893** |
| EC 3.1.3.1 | Sequence-only | 5,159 | 0.803 | 0.918 | 0.857 |
|  | System-aware | 5,159 | 0.803 | 0.918 | 0.857 |
| EC 3.5.1.5 | Sequence-only | 4,860 | **0.534** | 0.798 | **0.640** |
|  | System-aware | 4,860 | 0.358 | **0.897** | 0.512 |
| EC 3.2.1.23 | Sequence-only | 4,785 | 0.582 | 0.915 | 0.712 |
|  | System-aware | 4,785 | 0.582 | 0.915 | 0.712 |
| EC 3.2.1.21 | Sequence-only | 4,777 | 0.543 | 0.907 | 0.680 |
|  | System-aware | 4,777 | 0.543 | 0.907 | 0.680 |
| EC 3.2.1.22 | Sequence-only | 4,559 | 0.432 | 0.913 | 0.586 |
|  | System-aware | 4,559 | 0.432 | 0.913 | 0.586 |
| EC 3.4.11.1 | Sequence-only | 4,526 | **0.890** | 0.769 | 0.825 |
|  | System-aware | 4,526 | 0.875 | **0.948** | **0.910** |
| EC 3.2.1.31 | Sequence-only | 4,473 | 0.303 | 0.269 | 0.285 |
|  | System-aware | 4,473 | 0.303 | 0.269 | 0.285 |
| EC 3.2.1.52 | Sequence-only | 4,465 | 0.322 | 0.962 | 0.482 |
|  | System-aware | 4,465 | 0.322 | 0.962 | 0.482 |
| EC 3.2.1.20 | Sequence-only | 4,449 | **0.735** | 0.558 | 0.634 |
|  | System-aware | 4,449 | 0.664 | **0.759** | **0.708** |

**Sequence-only** indicates the performance of ML-derived scores. **System-aware** indicates the performance of refinement scores through genome-scale metabolic constraints. Bold values highlighting the superior performance in each category.

The optimization framework exhibited a targeted refinement effect, particularly for enzymes serving as metabolic bottlenecks.

For instance, *α*-glucosidase (EC 3.2.1.20) and aminopeptidase (EC 3.4.11.1) showed recall increases of 20.1% and 17.9%, respectively. These gains reflect the ability of system-aware constraints to recover enzymes that are topologically indispensable for maintaining steady-state flux through carbon and nitrogen utilization pathways. Conversely, performance metrics remained stable for enzymes that are not required for the biomass production objective or only active under certain conditions not represented by our experiment.

To ensure the structural integrity of this large-scale dataset, we used the MEMOTE benchmarking suite (Lieven *et al*., 2020). The reconstructed models achieved an average consistency score of approximately 0.70, reflecting high standards for stoichiometry and connectivity. Furthermore, Flux Balance Analysis (FBA) confirmed the physiological viability of the models, achieve a median growth rate of 7.4*h*^−1^ (Figure S4). These results suggest that METEOR recovers essential metabolic pathways that standard methods omit, resulting in a more authentic representation of phenotypic potential.

## Discussion

### Systemic constraints and effects on precision and recall

Our evaluation on the BacDive dataset reveals a divergence between sequence-based precision and system-based viability. While METEOR significantly improved the recovery of experimentally realized enzymatic activities (Recall: 0.751), we observed a decrease in precision. This shift here is likely not just a statistical trade-off but reflects the difference between a function annotation being present in a database or biologically necessary.

Standard metrics treat any prediction absent from the ground-truth database as a False Positive. However, biological databases are inherently incomplete. When METEOR predicts an enzyme that the baseline omits, it does so because that enzyme is topologically required to synthesize essential biomass components (e.g., amino acids, nucleotides). If the organism is known to grow in the simulated environment, these “False Positives” may represent unannotated but essential gene functions rather than computational errors. Consequently, the decline in precision likely results from incomplete annotations rather than incorrect predictions.

### Boundary conditions for metabolic refinement

The reliance on a global biomass objective imposes specific boundary conditions on the refinement process. METEOR is a constraint-based filter, not a universal correction tool. Consequently, there are distinct biological contexts where our method will effectively remain silent, never updating the probabilistic scores of an enzyme regardless of its presence in the genome.

The MILP formulation minimizes the cost to produce core biomass components. Enzymes involved solely in secondary metabolic pathways (Michal, 1999), such as those for antibiotic synthesis (Martin, 1998), pigment production (Hearing and Tsukamoto, 1991; Mohammadi *et al*., 2012), or bioluminescence (Brodl *et al*., 2018), do not contribute to this objective. Unless these metabolites are explicitly added to the biomass function, the algorithm perceives no systemic “need” for these enzymes and will not recruit them.

The optimization includes a parsimony term (*λ* ∑|*v*_*j*_ |) to favor minimal flux distributions. If a pathway is already satisfied by a high-confidence primary enzyme, the solver incurs a penalty for activating a lower-confidence isozyme. Consequently, redundant enzymes with weak sequence signals will remain unverified, as the system can function optimally without them.

Furthermore, our validation simulated a “complete” medium to maximize theoretical potential. However, enzymes required only under specific stress conditions (e.g., acid shock proteins) or for the uptake of exotic nutrients not present in the simulation environment will never carry essential flux. Without a direct link to growth in the simulated condition, the algorithm lacks the ability to refine these predictions.

Also, the algorithm weighs the cost of adding a reaction against the benefit of satisfying the biomass constraint. If a missing enzyme is part of a long, disconnected pathway requiring multiple gap-filling steps (high cumulative penalty) to connect to the core network, the optimizer may determine that the cost outweighs the benefit. In these cases, the entire pathway, including the potential enzyme, will remain dormant.

Finally, we restrict the current application of METEOR to prokaryotic systems. Eukaryotic metabolism, particularly in mammals, relies on strict intracellular compartmentalization to segregate conflicting thermodynamic processes (Martin, 2010). Because our current MILP formulation models the cell as a single well-mixed reaction volume, it lacks the specific transport constraints required to represent organelle-specific activity. Applying the current framework to eukaryotic genomes without accounting for subcellular localization would likely yield biologically infeasible flux solutions. Therefore, the extension of METEOR to higher organisms requires the integration of localization prediction signals into the constraint matrix.

### Structural evidence supports system-driven re-annotation

To illustrate how metabolic constraints can correct sequence-based protein WP 187575949.1 (UniParc UPI00165674F7) in *Shewanella* algae. The sequence-only baseline (DeepProZyme) annotated this protein as an endo-1,4-beta-xylanase (EC 3.2.1.8). However, METEOR, driven by the incompatibility of xylanase activity with the local metabolic network requirements, identified this as a potential misannotation and reassigned high probability to peptidase activities (EC 3.4.21.50/EC 3.4.24.28).

Orthogonal structural analysis supports this system-driven correction. Domain analysis via InterPro (Jones *et al*., 2014) confirms the presence of the AbiJ N-terminal domain (PF18863), a component of bacterial anti-phage defense systems typically associated with proteolytic rather than xylanolytic activity (Anantharaman *et al*., 2013). Furthermore, physicochemical profiling via DoGSite3 (Graef *et al*., 2023) of the predicted binding pocket reveals an extreme hydrophobicity (0.76) and a confined volume (193.02Å^3^). While xylanases require hydrophilic, open clefts to bind bulky polysaccharide chains (Pastor *et al*., 2007), these structural features are characteristic of peptidase active sites optimized for peptide bonds. This case demonstrates that METEOR effectively filters incorrect functional annotations generated by sequence classifiers, using global metabolic logic to enforce biologically plausible assignments even in the absence of direct structural inputs.

### Mechanisms of resilience in fragmented genomes

Our results demonstrate that METEOR can achieve high annotation correctness even when 90% of the genome is removed. This resilience is not due to the inherent scale-free topology of the network, as random removal of hub nodes typically collapses network connectivity. Rather, it arises from constraint-driven selection.

In the genome incompleteness experiments, the removal of “background” proteins drastically reduced the solution space. The Mixed-Integer Linear Programming (MILP) optimizer was forced to rely on the remaining high-potential candidates (the test set) to satisfy the mandatory biomass objective. The systemic necessity of these proteins became absolute because no alternative candidates remained to fulfill their metabolic roles. This finding implies that even in highly fragmented metagenome-assembled genomes (MAGs), METEOR can recover essential functional annotations by anchoring predictions to the immutable requirements of biomass synthesis, effectively using the metabolic model as a scaffold to organize sparse genomic data.

## Conclusion

We developed METEOR, a framework that integrates continuous enzyme prediction scores from deep learning models directly into genome-scale metabolic reconstruction. By embedding enzymatic evidence within a Mixed-Integer Linear Programming formulation, our approach refines functional annotation through global metabolic feasibility rather than treating it as an isolated, protein-level task. Focusing on microbial genomes, we demonstrated that enforcing system-level constraints enhances both the functional coherence and physiological viability of reconstructed metabolic networks across diverse bacterial species.

Contemporary machine learning strategies for protein annotation have largely adhered to a reductionist paradigm (Tawfiq et al., 2025), optimizing predictive accuracy on individual sequences while ignoring the cellular context in which these molecules operate. Our results highlight the limitations of this component-centric view. We found that network-level consistency, specifically the requirement to satisfy biomass production, resolves ambiguities that local sequence features cannot address. Just as the emergence of systems biology in the early 2000s (Kitano, 2002) shifted the analytical focus from isolated parts to emergent interactions, the next generation of biological machine learning must move beyond independent classification to model the thermodynamic and stoichiometric logic that governs life.

METEOR represents a step toward this unified inference problem demonstrating that global system constraints act as a necessary form of biological supervision for data-driven predictions. Future work will extend this framework to incorporate condition-specific objectives and alternative metabolic tasks, further establishing that the accurate annotation of the genotype requires a quantitative understanding of the phenotype.

## Acknowledgements

We acknowledge support from the KAUST Supercomputing Laboratory.

