## Supplementary material for "METEOR: joint genome-scale reconstruction and enzyme prediction": Supplementary.pdf

### S1 Supplementary Tables

Detailed information of the 22 bacterial genomes used in the Price-149 dataset evaluation.This table includes assembly accessions, genome sizes, and the number of proteins per genome. (Attached file `Table1_Price_genome_metainfo.csv` )

### S2 Supplementary Figures

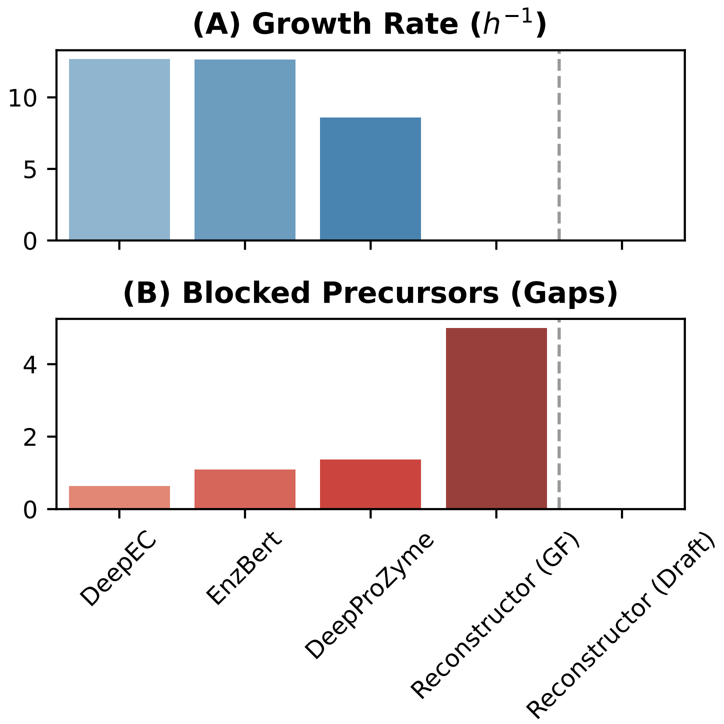


Figure S1: **omparative analysis of predicted growth rates and model integrity.** Models refined by METEOR across all underlying baselines achieved robust growth (median $> 7.4\,h^{-1}$, confirming their readiness for flux balance analysis. Conversely, Reconstructor-derived models failed to sustain metabolic flux under default medium conditions and exhibited substantial pathway gaps.

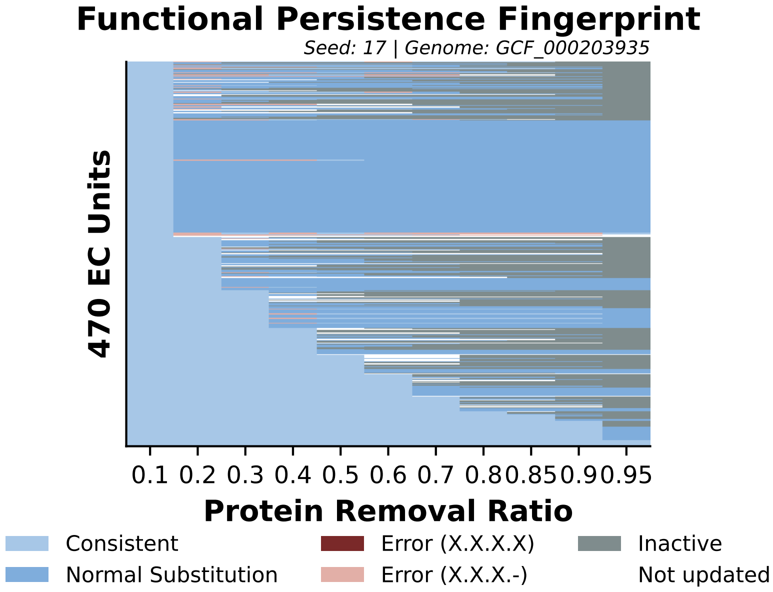


Figure S2: **Retention of EC–Protein Associations under Increasing Genome Removal Ratios.** Elucidates this functional persistence behavior through a single-genome case study from *Dinoroseobacter shibae*. The visualization demonstrates that our method favors “Consistent (Light Blue)” updates, where the most potential candidate protein is persistently selected for same EC unit across varying levels of genome fragmentation. “Normal Substitution (Blue)” occurs only when the primary candidate is removed, prompting the optimizer to shift the assignment to the next best available candidate.

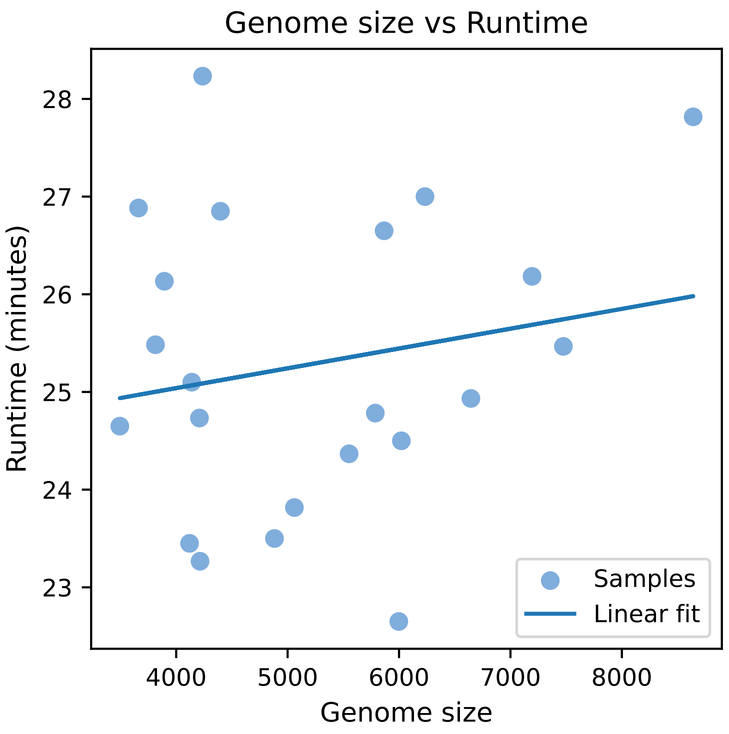


Figure S3: **Computational scalability analysis.** Total execution time (minutes) is plotted against genome size (number of proteins), exhibiting a near-linear scaling relationship. The efficiency of the evidence-weighted MILP formulation allows for the joint reconstruction and refinement of a standard bacterial proteome in approximately 25 minutes, with even large-scale genomes (∼9,000 proteins) processed in under 30 minutes. This performance underscores the framework's suitability for high-throughput genomic and metagenomic annotation pipelines.

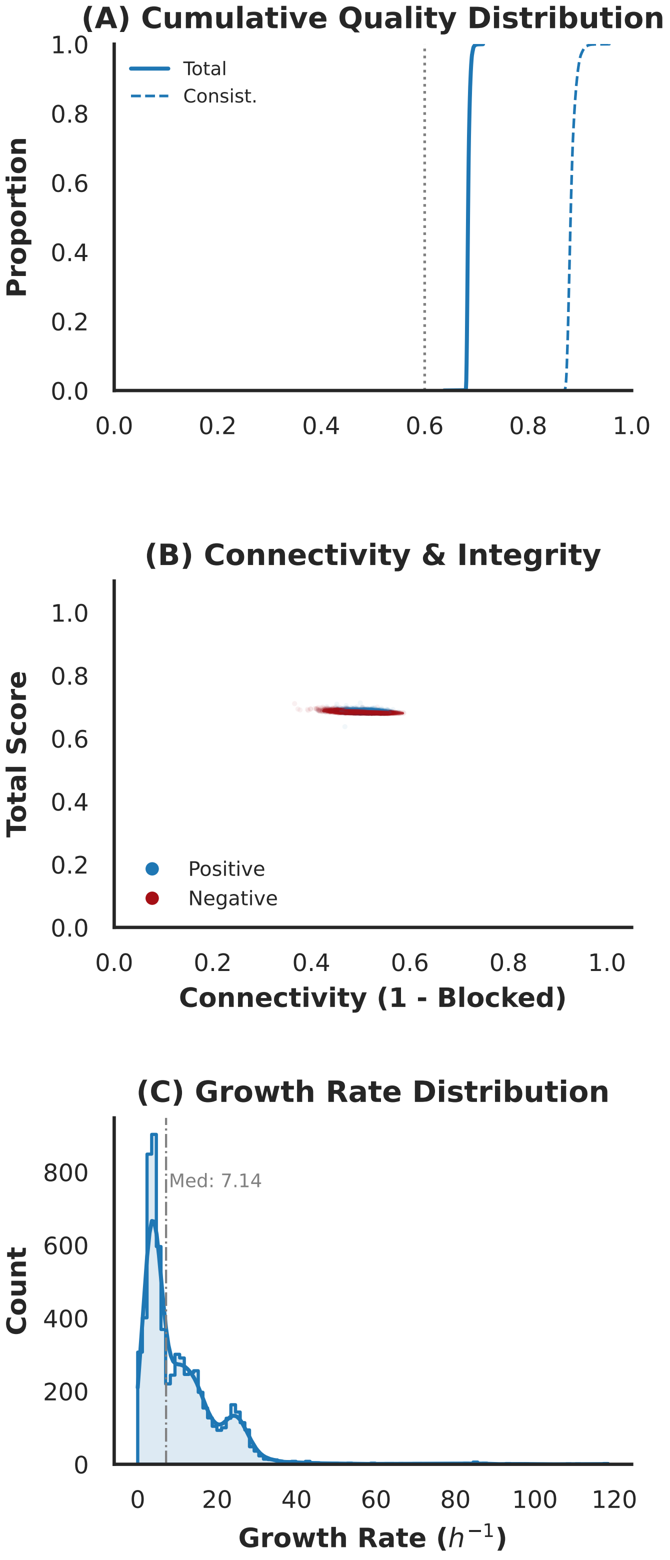


Figure S4: **Comprehensive quality evaluation of metabolic models across 6,894 BacDive genomes.** (A) Cumulative distribution of MEMOTE quality metrics, illustrating that the majority of reconstructed models attain a Total score of 0.7 and a stoichiometric Consistency score of 0.9. (B) Topological characterization (Connectivity and Integrity) across the dataset, demonstrating comparable network robustness between Gram-positive and Gram-negative bacteria. (C) Statistical distribution of predicted growth rates ($h^{-1}$); the METEOR-derived models exhibit robust physiological viability with a median growth rate of 7.14 $h^{-1}$, confirming the successful restoration of essential biomass synthesis pathways.

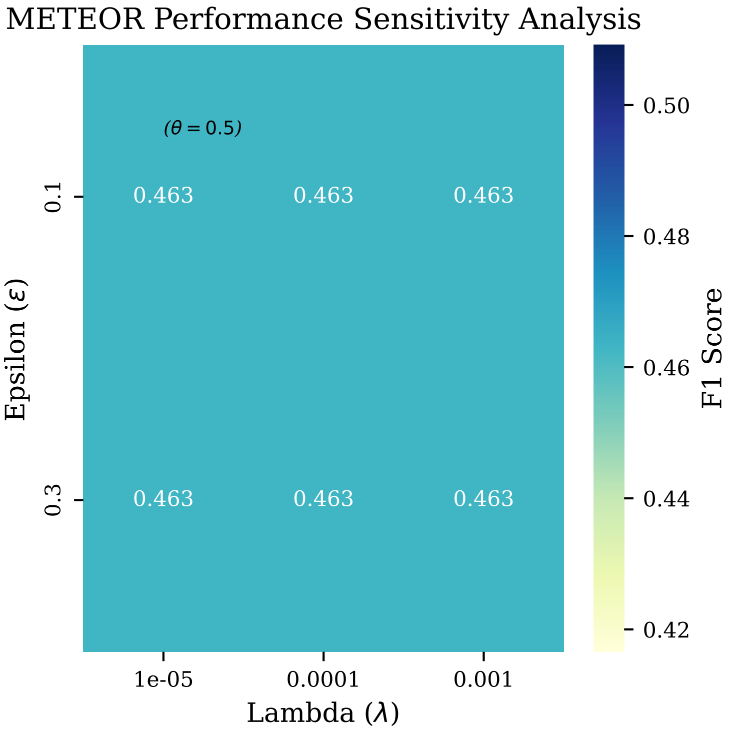


Figure S5: **Comprehensive evaluation of hyperparameters of $\epsilon$ and $\lambda$.** METEORis not overly dependent on hyperparameters tuning.
